# Shugoshin holds the potential to inhibit APC/C and thereby prevents separase activation

**DOI:** 10.1101/2025.09.22.677724

**Authors:** Ke Zhang, Shuchen Guo, Li Sun, Haitong Hou, Yoshinori Watanabe

## Abstract

Accurate chromosome segregation during mitosis and meiosis depends on shugoshin protein which localizes at the centromere of chromosomes. Sgo1, a meiosis-specific shugoshin in fission yeast, collaborates with the PP2A phosphatase enzyme to remove phosphate groups from Rec8 cohesin, thereby preventing its cleavage by separase during meiosis. Here we show that ectopic expression of Sgo1 causes chromosome segregation errors during mitosis, especially in *cut1-206* separase mutant cells, but that the D-box mutant Sgo1 has no effect in *cut1-206* cells. Our molecular genetic analyses suggest that the D-box of Sgo1 directly binds Slp1/CDC20 and inhibits APC/C-dependent separase activation, thereby preventing cleavage of Rad21 cohesin. In contrast, during meiosis, the Sgo1 D-box mutant makes the *cut1-206* mutant more difficult to cleave Rec8 cohesin. This is because the D-box mutation increases the amount of Sgo1 protein during meiosis, thereby enhancing its canonical protective ability against Rec8 cohesin. Thus, our studies using a separase mutant have revealed that Sgo1 has multiple faces to protect coheisn.

## Introduction

In proliferating fission yeast cells, duplicated chromosomes (sister chromatids) become connected during S phase through the action of the cohesin complex, which consists of four core subunits: two SMC (structural maintenance of chromosome) family ATPase proteins, Psm1 and Psm3, a kleisin family protein Rad21, and accessory subunit Psc3 (Tomonaga et al. 2000). The cohesion of sister chromatids is maintained throughout G2 phase until metaphase when chromosomes are aligned at the spindle equator. When anaphase begins, the anaphase-promoting complex/cyclosome (APC/C) triggers the degradation of securin, a protein that keeps separase inactive. Once activated, separase cleaves the kleisin subunit Rad21, opening the cohesin ring and separating the sister chromatids (Yanagida 2000; Peters et al. 2008; Nasmyth and Haering 2009). APC/C activity is tightly controlled on different levels. Several APC/C subunits are known to be phosphorylated by Cdks and polo-like kinase during mitosis (Kraft et al. 2003; Pines 2011). In addition to phosphorylation, APC/C activity is controlled by the spindle assembly checkpoint (SAC), which is activated by the absence of tension or lack of microtubule occupancy at the kinetochore (Musacchio and Salmon 2007; McAinsh and Kops 2023). Consequently, separase is activated only when all chromosomes are correctly attached to the spindle in a bipolar orientation, ensuring accurate chromosome separation during anaphase.

In meiosis, Rad21 subunits are largely replaced by the meiosis-specific Rec8, a key protein that establishes meiosis-specific chromosome segregation, resulting in the production of four haploid nuclei, or gametes (Watanabe and Nurse 1999). During the meiosis I division, sister kinetochores are captured by microtubules from the same pole (mono-orientation) and Rec8 is cleaved by separase along the chromosome arms but not at the centromere (cohesion protection) (Buonomo et al. 2000; Kitajima et al. 2003). Shugoshin (Sgo1) and PP2A collaboratively antagonize casein kinase 1 (CK1)-dependent Rec8 phosphorylation, a prerequisite for cleavage by separase, thus protecting Rec8 cohesin from cleavage at the centromere (Kitajima et al. 2004; Kitajima et al. 2006; Riedel et al. 2006; Ishiguro et al. 2010; Katis et al. 2010). The B55 and B56 regulatory subunits of PP2A are fully conserved from yeasts to humans, acting as major determinants of PP2A holoenzyme specificity. PP2A-B56 is generally used to protect cohesin by shugoshin during meiosis (Kitajima et al. 2006; Riedel et al. 2006; Lee et al. 2008). On the other hand, PP2A-B55 is known to play a role in attenuating the cleavage of cohesin in proliferating cells of budding yeast (Clift et al. 2009; Yaakov et al. 2012).

Like fission yeast Sgo1, mouse SGO2 (shugoshin 2) and its associated PP2A play a key role in protecting REC8 cohesin from separase cleavage during meiosis in germ cells (Lee et al. 2008; Llano et al. 2008; El Yakoubi et al. 2017). However, recent studies suggest that SGO2 binds to MAD2 when the SAC is activated, and the complex blocks the active site of separase, preventing the cleavage of Rad21 cohesin in somatic cells (Orth et al. 2011; Hellmuth et al. 2020). In this study, we analyzed whether Sgo1 in fission yeast also has the ability to inhibit separase activity in proliferating cells.

## Results

### Ectopic expression of Sgo1 causes chromosome segregation errors during mitosis

Although Sgo1 is normally expressed only during meiosis, ectopic expression of Sgo1 in proliferating cells under the strong promoter Pnmt1 (but not the weak promoter Pnmt41) leads to chromosome nondisjunction only if Rec8 cohesin is co-expressed in the cells (Kitajima et al. 2004) (Fig. 1A). This indicates that the cleavage of Rec8 but not Rad21 is prevented by Sgo1 in proliferating cells. Notably, however, the ectopic expression of Sgo1 causes lethality in the mutant of separase (*cut1-206*) despite the absence of Rec8 in the cells (Kitajima et al. 2004) (Fig. 1B). Sgo1-L299A protein which has a mutation in the SGO domain and fails to localize at centromeres (Fig. EV1A,B), did not affect survival of *cut1-206* cells (Fig. 1B). These results suggested that centromere localization of Sgo1 is a key feature that affects survival of *cut1-206* cells.

**Figure 1.**
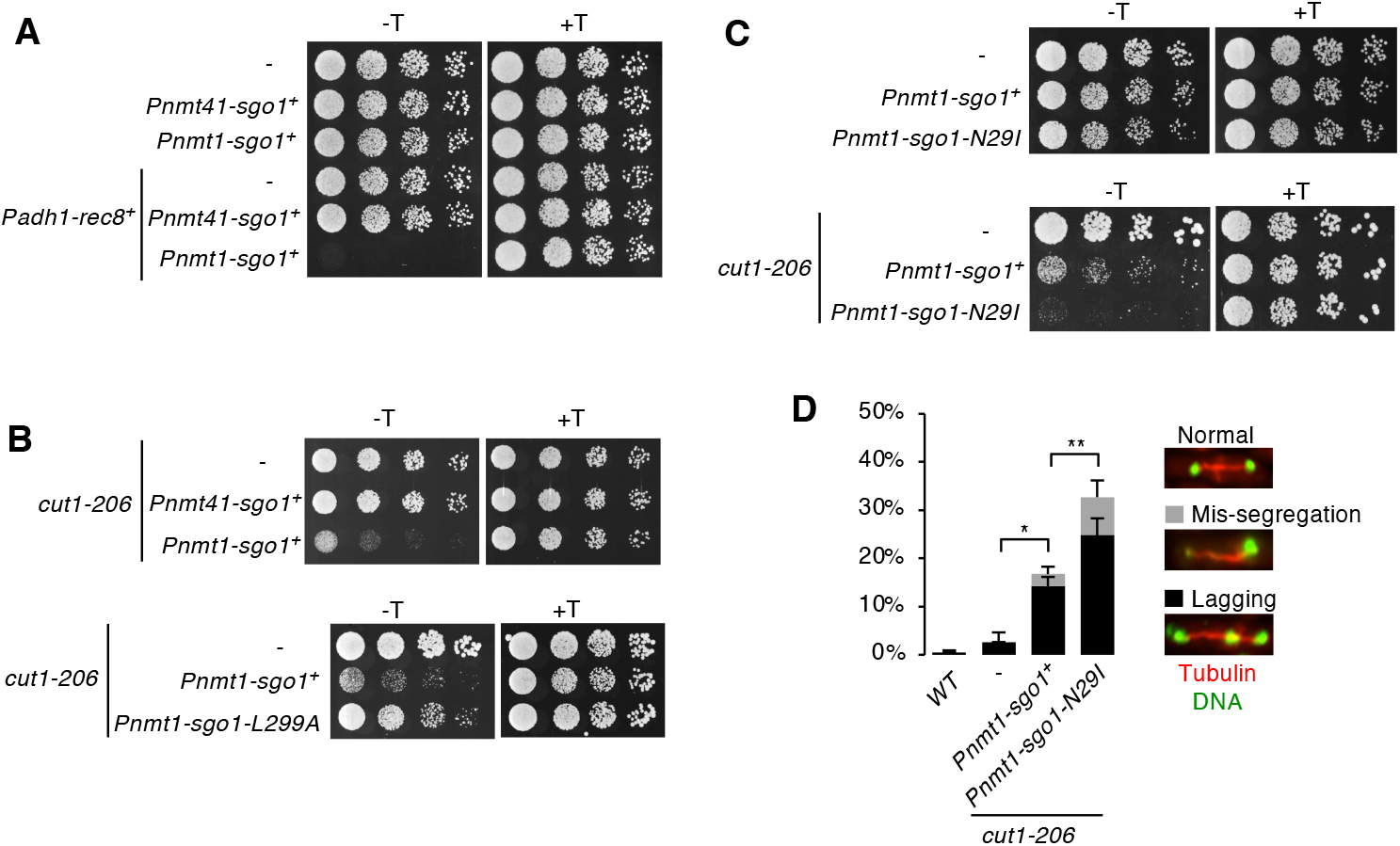
Ectopic expression of Sgo1 affects survival of *cut1-206* cells. **A, B, C**, Serial 5-fold dilutions of the indicated cells were spotted on EMM plates including (+T) or lacking (-T) thiamine and grown at 30°C; *cut1-206* cells were grown at 28°C. The absence of thiamine induces expression from the Pnmt1 and Pnmt41 promoters. **D**, The indicated cells were stained for DNA. The spindles were visualized by expressing mCherry-Atb2 (α2-tubulin). Frequencies of lagging chromosomes and mis-segregation at anaphase cells. Examples are shown at the right. Error bars, SD (three independent experiments, n > 100 anaphase cells in each experiment); \**P* < 0.05, \*\**P* < 0.01.

Because Sgo1 utilizes PP2A to prevent separase-mediated cleavage of Rec8 cohesin, we investigated whether chromosome mis-segregation caused by Sgo1 in *cut1-206* cells was also due to PP2A bound to Sgo1. For that purpose, we made a mutant in the coiled-coil domain of Sgo1 (Sgo1-N29I) that fails to recruit PP2A and therefore cannot protect Rec8 cohesin during meiosis (Fig. EV1C-F). Surprisingly, ectopic expression of Sgo1-N29I increased, rather than decreased, lethality *in cut1-206* cells compared to wild-type Sgo1 (Fig. 1C). Nuclear staining of anaphase cells showed that ectopic expression of Sgo1 and Sgo1-N29I enhanced chromosome segregation errors in *cut1-206* mutant cells, often exhibiting lagging chromosomes and mis-segregation of chromosomes at anaphase (Fig. 1D). These results suggest that ectopically expressed Sgo1 weakened separase activity, preventing separase-mediated cleavage of Rad21, independently of PP2A.

A recent study showed that mammalian SGO2 binds to the conserved separase catalytic site through a noncleavable pseudosubstrate sequence in SGO2 (φExxX, where φ denotes a hydrophobic residue, x denotes any residue, and X denotes any residue other than R), revealing that mammalian SGO2, similar to securin, inhibits separase (Hellmuth et al. 2020). To examine the possibility that fission yeast Sgo1 inhibits separase through a similar mechanism, four putative pseudo-substrate sequences of Sgo1 were identified and the glutamic acid residues within these sequences were replaced with alanine. As a result, the mutant Sgo1 proteins affected survival of *cut1-206* cells similarly to wild-type Sgo1 (Fig. EV2A,B), implying that these putative pseudo-substrate sequences do not contribute to separase inhibition and that the mechanism of separase inhibition by fission yeast Sgo1 is distinct from that of mammalian SGO2.

### The Sgo1 D-box mutant fails to affect survival of *cut1-206* cells

Given that APC/C activity is essential for separase activation, we sought a possibility that Sgo1 might inhibit APC/C rather than separase. As reported previously (Kitajima et al. 2004), Sgo1 is degraded during anaphase I in part depending on Slp1 (Cdc20 homolog), a substrate-recognition subunit and activator of the APC/C. Aligning the D-box sequences (RxxLxxxN) of various proteins revealed that Sgo1 from fission yeast, budding yeast, and humans shares non-canonical D-box sequence (XxxLxxxN, X is not R)(Karamysheva et al. 2009; Eshleman and Morgan 2014). We made a mutant Sgo1-L185A in which the conserved leucine amino acid in the D-box like sequence was replaced by alanine (Fig. 2A). We found the L185 is indeed required for timely degradation of Sgo1 during anaphase I because Sgo1-L185A proteins persisted at the centromeres beyond meiosis I until meiosis II (Fig. EV2C,D). Two hybrid assays indicated that Sgo1 binds Slp1 and the binding ability was reduced in Sgo1-L185A protein (Fig. 2B). Surprisingly, in contrast to wild-type Sgo1, Sgo1-L185A did not significantly affect survival of *cut1-206* cells (Fig. 2C), suggesting that the D-box of Sgo1, which can bind to Slp1, plays a key role in affecting survival of *cut1-206* cells. It is also known that the spindle assembly checkpoint protein Mad2 binds to Slp1 to inhibit APC/C (May and Hardwick 2006). Indeed, overexpression of Mad2 and Sgo1 synergized in proliferating cells, increasing chromosome segregation errors during mitosis, with Sgo1-L185A showing a lesser effect (Fig. 2D,E). These results collectively support the idea that ectopically expressed Sgo1 inhibits APC/C through the interaction of the Sgo1 D-box with Slp1, an activator of APC/C.

**Figure 2.**
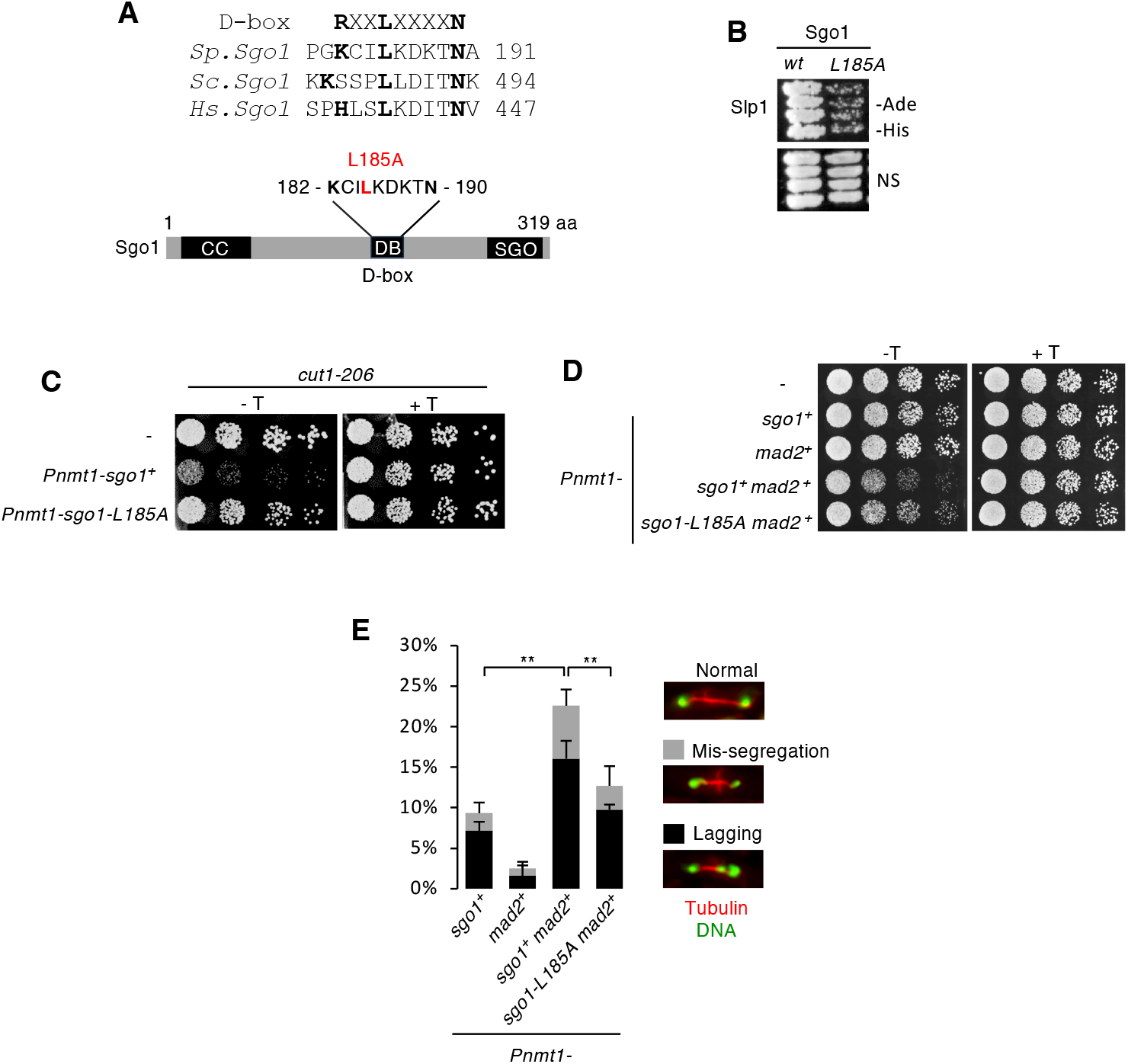
The Sgo1 D-box mutant fails to affect survival of *cut1-206* cells. **A**, Alignment of the D-box motif in Sgo1of *S. pombe, S. cerevisiae* and *H. sapiens*. Schematic diagram of the Sgo1 protein showing the PP2A-interacting coiled-coil region (CC), D-box (DB), and SGO motif (SGO). D-box sequence and mutation site are shown. **B**, Yeast two-hybrid assay testing the interaction of Slp1 and Sgo1. The transformants were grown on a nonselective plate and plate lacking adenine and histidine. **C**, Serial 5-fold dilutions of the indicated cells were spotted on EMM plates including (+T) or lacking (-T) thiamine and grown at 28°C. **D**, Serial 5-fold dilutions of the indicated cells were spotted on EMM plates including or lacking thiamine and grown at 30°C. **E**, The indicated cells were stained for DNA. The spindles were visualized by expressing mCherry-Atb2 (α2-tubulin). Frequencies of lagging chromosomes and mis-segregation at anaphase cells. Examples are shown at the right. Error bars, SD (three independent experiments, n > 100 anaphase cells in each experiment); \*\**P* < 0.01.

### Phosphorylation of the D-box of Sgo1 may enhance its inhibitory function against APC/C

Of note, yeast and human Sgo1 bears a polo-like kinase consensus sequence (D/E x S/T) within the D-box sequence (Alfieri et al. 2017) (Fig. 3A). This sequence also exists in the non-canonical D-box sequence (RxxLxxxI) of fission yeast Mes1, a meiosis-specific competitive substrate for Slp1 (Kimata et al. 2008) (Fig. 3A). To investigate the role of this potential phosphorylation at T189 of Sgo1, we introduced the non-phosphorylatable T189A and phosphomimetic T189E mutations at T189 of Sgo1 and examined the effect of the mutations on the survival of *cut1-206* cells. In contrast to Sgo1-WT, Sgo1-T189A did not affect survival of *cut1-206* cells, whereas Sgo1-T189E had an effect equal to or greater than that of Sgo1-WT (Fig. 3B). In yeast two-hybrid assays, Sgo1-T189A bound less strongly to Slp1 than Sgo1-WT, whereas Sgo1-T189E bound well to Slp1 (Fig. 3C). These results are consistent with the hypothesis that Sgo1-Slp1 interaction is regulated by phosphorylation at T189 in the D-box sequence of Sgo1. Because the effect of Sgo1-N29I was similar to Sgo1-T189E and stronger than Sgo1-T189A, we speculate that phosphorylation at T189 is negatively regulated by PP2A bound to Sgo1 (Fig. 3D).

**Figure 3.**
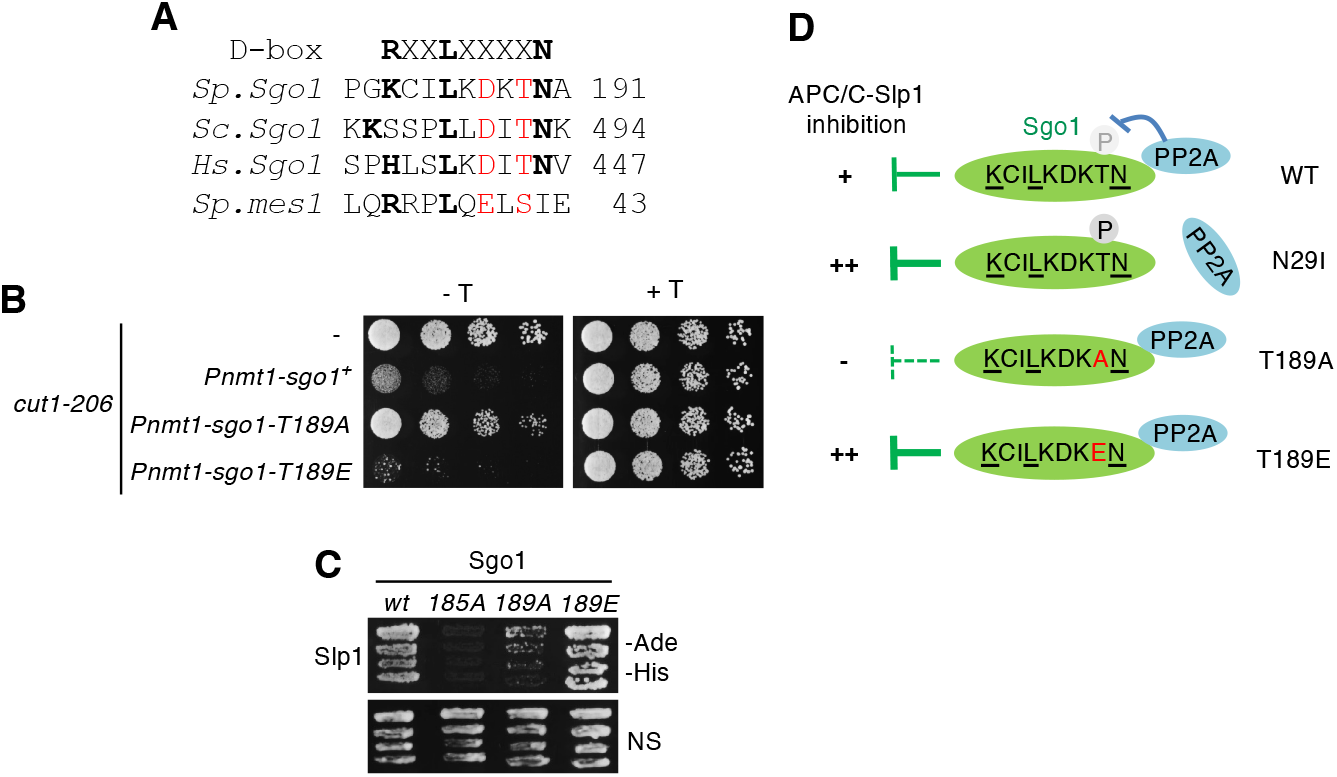
Phosphorylation in the D-box of Sgo1 may enhance its inhibition of the APC/C. **A**, Alignment of the D-box motif in Sgo1. The D-box motif of Mes1 of *S. pombe* is also shown. The Polo-like kinase consensus sequence (D/E x S/T) is shown in red. **B**, Serial 5-fold dilutions of the indicated cells were spotted on EMM plates including (+T) or lacking (-T) thiamine and grown at 28°C. **C**, Yeast two-hybrid assay testing the interaction of Slp1 with Sgo1 (amino acid 81-211) and its mutants. The transformants were grown on the nonselective plate and a plate lacking adenine and histidine. **D**, Schematic diagram of the inhibitory effect of Sgo1 and its D-box mutants on APC/C-Slp1.

### PP2A-B56 and D-box-independent protection of Rec8 by Sgo1 in proliferating cells

To further explore the potential of ectopically expressed Sgo1, we investigated how Sgo1 protects Rec8 in mitotic cells compared with meiotic cells. We examined meiotic chromosome segregation by monitoring GFP fluorescence associated with a *lacO* array integrated at the centromere (*imr1*::GFP). During meiosis, Sgo1 prevents the cleavage of Rec8 through Sgo1-associated PP2A and Rec8 is not protected in cells lacking the PP2A regulatory B56 subunits, Par1 and Par2 (Kitajima et al. 2006) (Fig. 4A,B). However, during mitosis, ectopic expression of Sgo1 caused lethality in *par1Δ par2*Δ cells only when Rec8 was co-expressed, indicating that Sgo1 can protect Rec8 independently of Par1 and Par2 (Fig. 4C). Accordingly, ectopic expression of Sgo1-N29I, which loses the interaction with the PP2A-B56 complex (containing Par1 or Par2), caused severe growth retardation in cells expressing Rec8. Remarkably, unlike the results in *cut1-206* cells, Sgo1-L185A caused lethality in cells expressing Rec8 just as Sgo1-WT did, suggesting that Sgo1 protects Rec8 completely independently of the APC/C-Slp1 inhibitory pathway (Fig. 4C). Another mutant Sgo1-L299A, which cannot localize to centromeres and loses the protective function in meiosis, did not cause lethality in Rec8 expressing cells (Fig. 4B,C). Taken together, these results suggest that centromere localization of Sgo1 is a key feature that allows it to protect Rec8 cohesin from separase-mediated cleavage in both mitosis and meiosis.

**Figure 4.**
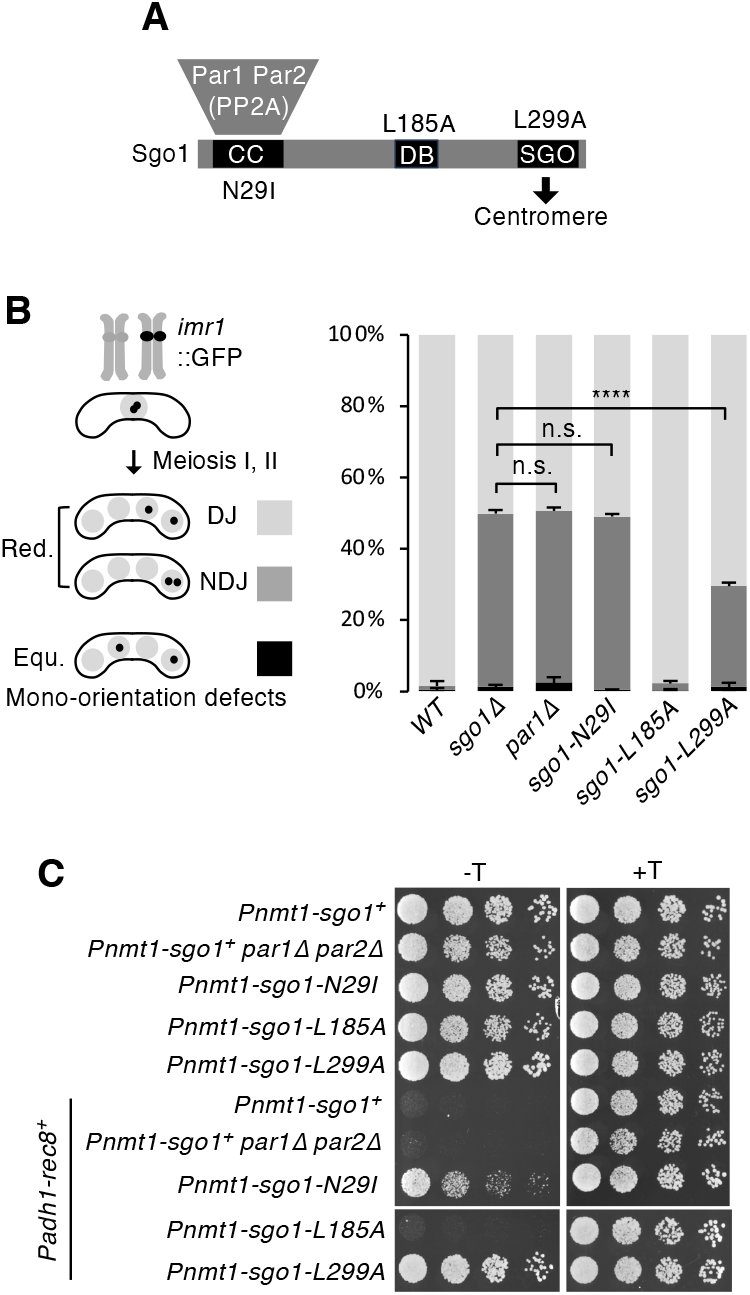
PP2A- and D-box-independent protection of Rec8 by Sgo1 in proliferating cells. **A**, Schematic diagram of the Sgo1 protein showing the PP2A-interacting coiled-coil region (CC), D-box (DB), and SGO motif (SGO), each containing mutation sites at N29, L185 and L299, respectively. **B**, Segregation pattern of *imr1*::GFP marked on one homolog in the indicated cells was monitored in cells after the meiosis II division. DJ: disjunction, meaning proper sister chromatid segregation in meiosis II. NDJ: nondisjunction, meaning sister chromatids move to the same pole in meiosis II, the defect originated from loss of cohesion during anaphase I. Red. (reductional): sister chromatids moved to the same pole in meiosis I. Equ. (equational): sister chromatids moved to opposite poles already in meiosis I. Error bars represent SD (three independent experiments, n > 150 cells in each experiment); \*\*\*\**P* < 0.0001. **C**, Serial 5-fold dilutions of the indicated cells were spotted on EMM plates including (+T) or lacking (-T) thiamine and grown at 30°C.

### The Sgo1 D-box mutant enhances the protection of Rec8 cohesin in meiosis

Although wild-type Sgo1 appeared at centromeres from prophase I to metaphase I and disappeared at anaphase I, Sgo1-L185A persisted at the centromere throughout meiosis (Fig. EV2C). Notably, compared with Sgo1-WT, the amount of Sgo1-L185A protein at centromeres during metaphase I was significantly increased (Fig. EV2D), suggesting that the level of Sgo1 protein during metaphase I is regulated not only at the synthesis level but also by APC/C-dependent degradation. In *sgo1-L185* cells, meiotic chromosome segregation was largely intact (Fig. 4B), indicating that the timely degradation of Sgo1 at anaphase I and its absence in meiosis II is not important for proper meiotic chromosome segregation. Because Sgo1 showed genetic interaction with Mad2 in proliferating cells (Fig. 2D), we investigated whether *sgo1-L185A* mutation affects SAC activation during meiosis. We measured the time from prometaphase I to anaphase I by monitoring the separation of SPBs (spindle pole bodies; initiation of spindle formation) and degradation of securin Cut2 (onset of anaphase I). In addition to wild-type cells, we examined *rec12Δ* cells, in which chromosomes are not properly aligned during metaphase I due to a lack of chiasmata between homologous chromosomes. Accordingly, the duration was 20 min in wild type cells, but was extended to 42 min in *rec12Δ* cells by SAC activation, whereas the *sgo1-L185A* mutation did not alter either duration (Fig. EV3A,B). These results suggest that Sgo1-dependent regulation of APC/C activity is absent in meiosis I.

To elucidate the genetic interaction between *cut1-206* and *sgo1-L185A* in meiosis, we analyzed chromosome segregation errors during meiosis I. To this end, we arrested cells after first meiotic division using the *mes1*-*B44* mutation. In response to elevated temperature, *cut1-206* cells showed an increase of nondisjunction of homologs during meiosis I, suggesting that reduced separase activity inhibited cleavage of Rec8 along the chromosomes (Fig 5A). Notably, at the semi-permissive temperature of 25°C, the frequency of nondisjunction of homologs was increased in *cut1-206 sgo1-L185A* cells compared to *cut1-206 sgo1*^*+*^ or *cut1-206 sgo1Δ* cells (Fig 5B). These results suggest that, in contract to the outcomes of mitosis (Fig. 2C), Sgo1-L185A has an enhanced ability to prevent cohesin cleavage by separase during meiosis I compared to wild-type Sgo1.

**Figure 5.**
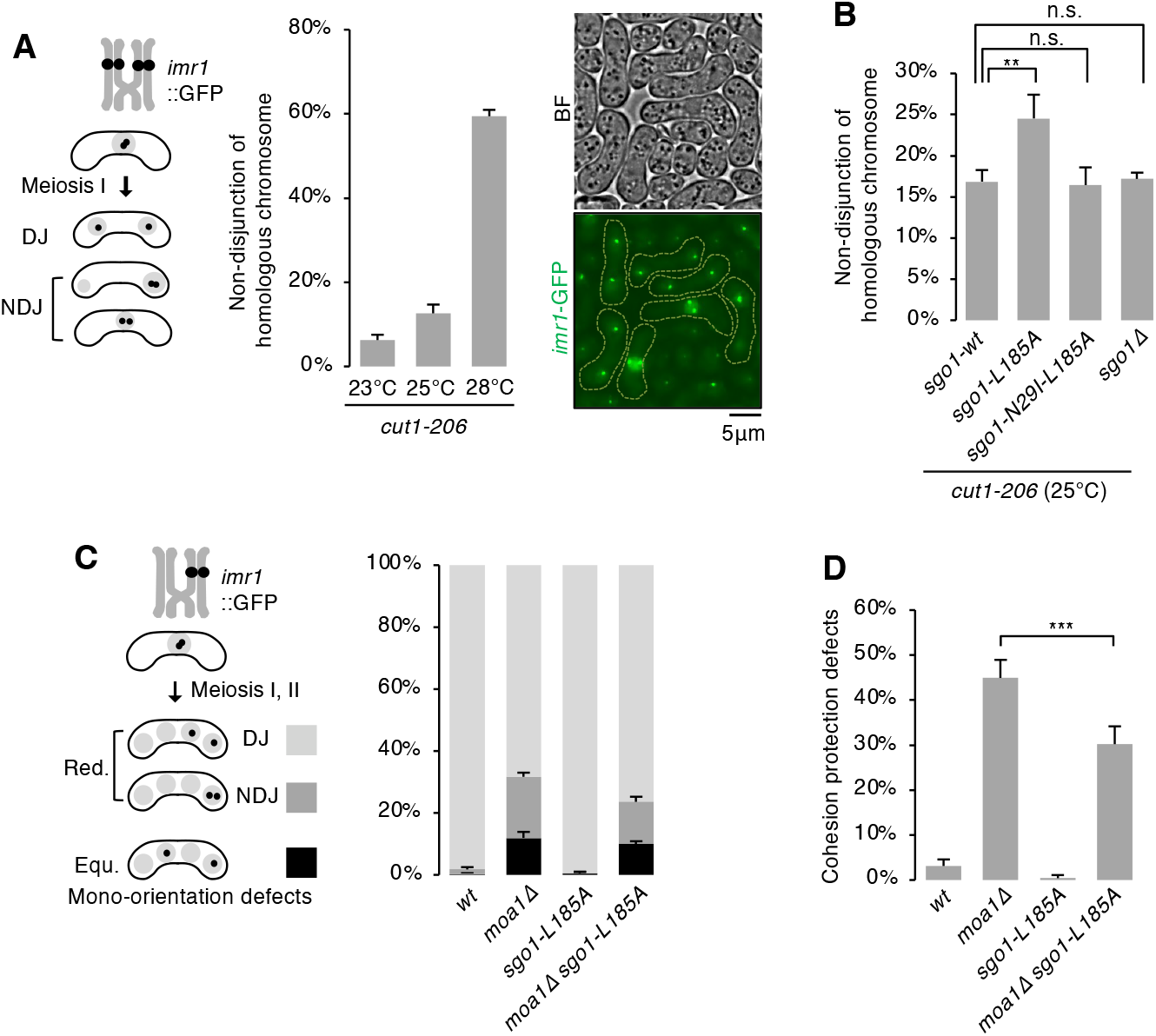
The Sgo1 D-box mutant enhances the protection of Rec8 in meiosis. **A**, Segregation pattern of *imr1*::GFP marked on both homologs was monitored in *cut1-206* cells cultured at 23°C, 25°C and 28°C and arrested at prophase II by *mes1-B44*. Representative brightfield (BF) and fluorescent images of *cut1-206* cells cultured at 25°C are shown. **B**, Segregation pattern of *imr1*::GFP marked on both homologs was monitored in the indicated cells cultured at 25°C and arrested at prophase II by *mes1-B44*. Error bars represent SD (three independent experiments, n > 150 cells in each experiment). n.s., not significant; \*\**P* < 0.01. **C**, Segregation pattern of *imr1*::GFP marked on one homolog was monitored in the indicated cells after the meiosis II division. **D**, Cohesion protection defects were calculated from the results in **C** by the formula (NDJ x 2 / DJ + NDJ) x 100 %. Error bars represent SD (three independent experiments, n > 150 cells in each experiment). n.s., not significant; \*\*\**P* < 0.01.

Because nondisjunction did not increase in *cut1-206 sgo1-N29I-L185A* cells, the protection ability of Sgo1-L185A largely depended on its bound PP2A (Fig. 5B). Because *moa1Δ* cells show partial defects in cohesion protection at meiosis I, we examined the possibility that *sgo1-L185A* suppresses the protection defects of *moa1Δ*. In *moa1Δ* cells, although a minority population of cells (12%) underwent equational segregation at meiosis I (because of defects in mono-orientation), the remaining majority (88%) underwent reductional segregation due to the presence of chiasmata and tension exerted across homologs. However, because *moa1Δ* cells are partially defective in cohesion protection (Miyazaki et al. 2017), 22.5% of the reductional population of *moa1Δ* cells showed NDJ in meiosis II (Fig. 5C), suggesting that 45% of the reductional population lost the cohesion and underwent random segregation in meiosis II (Fig. 5D). Interestingly, this cohesion loss was reduced into 30% by *sgo1-L185A* (Fig. 5D). Together, these results suggest that Sgo1-L185A has enhanced protective capacity during meiosis as opposed to mitosis. In conclusion, ectopic expression of Sgo1 inhibits APC/C-separase activation axis in proliferating cells, but this function is apparently silent during meiosis, where Sgo1-bound PP2A instead plays a key role in preventing cohesin cleavage.

## Discussion

Fission yeast possess two shugoshin-like proteins, Sgo1 and Sgo2, both of which localize to the pericentromeric heterochromatin region, the site enriched with cohesin (Kitajima et al. 2004). Sgo1 is meiosis I-specific and binds to PP2A-B56, protecting centromeric Rec8 cohesin from separase-mediated cleavage during meiosis I (Kitajima et al. 2006; Riedel et al. 2006). Sgo2 is expressed in both mitosis and meiosis, and plays a role in establishing proper attachment of kinetochores to microtubules by recruiting Aurora B kinase to the centromere (Kawashima et al. 2007). On the other hand, budding yeast possess only one shugoshin protein Sgo1, which carries at least two functions; cohesion protection only in meiosis and Aurora B recruitment in both mitosis and meiosis (Katis et al. 2004; Marston 2014). Thus, it has been long believed that shugoshin does not have a function for cohesion protection in proliferating cells in yeasts. In proliferating human cells, SGO1-bound PP2A dephosphorylates the cohesin cofactors SA2 and sororin, preventing cohesin release during mitotic prophase (Hauf et al. 2005; Nishiyama et al. 2013). Human SGO1 also competes with the cohesin release factor Wpl for cohesin binding, thereby strengthening protection against cohesin release (Hara et al. 2014). Mammalian SGO2 can inhibit separase similarly to securin in proliferating cells (Hellmuth et al. 2020). Thus, shugoshin has multiple face to protect cohesin in eukaryotic cells (Fig. 6A).

**Figure 6.**
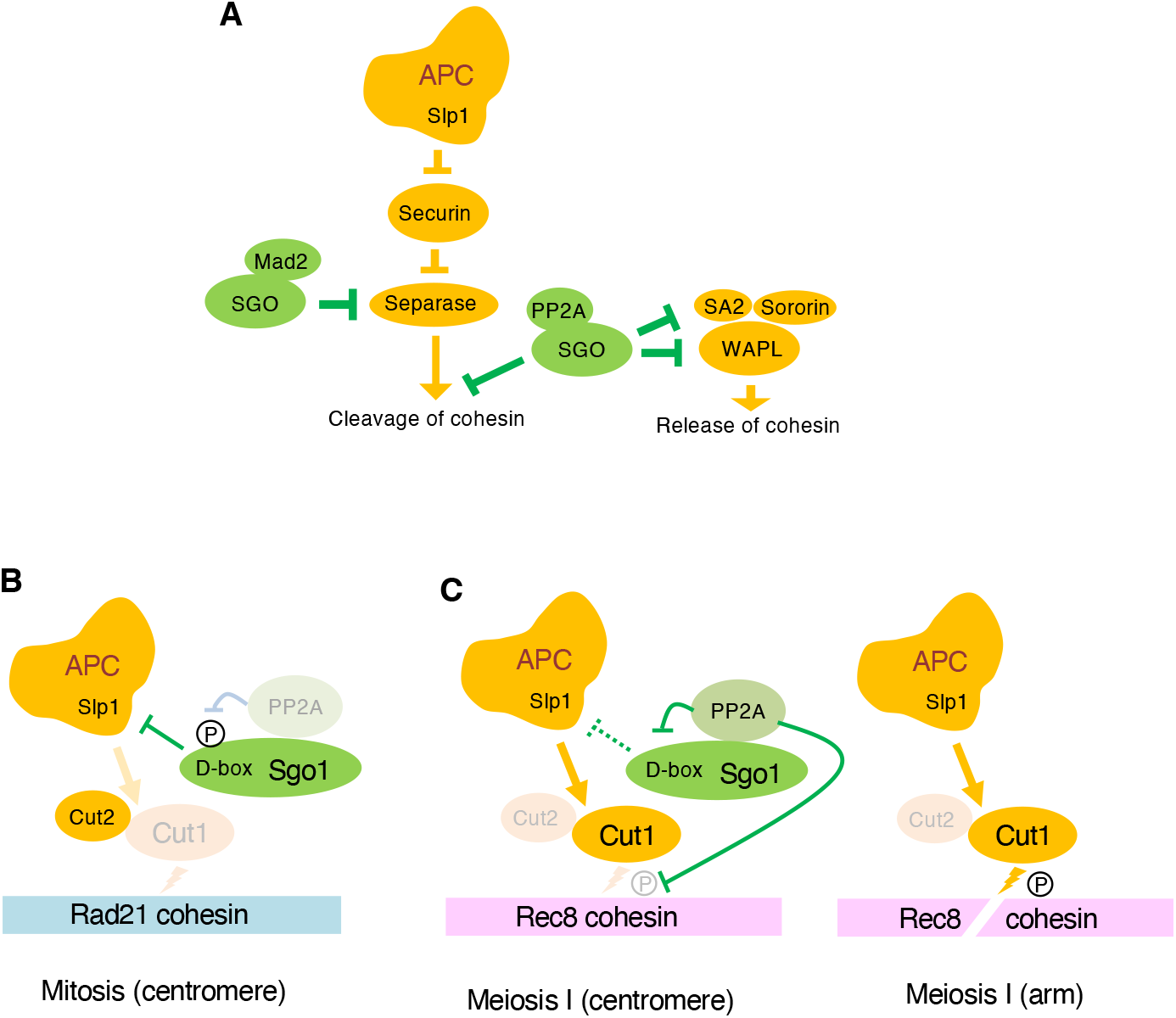
Schematic model of Sgo1 dependent protection of cohesin in mitosis and meiosis. **A**, Schematic diagram of the multiple faces of shugoshin (SGO) protecting cohesin in eukaryotic cells. **B**, Phosphorylation of the Sgo1-D-box promotes Sgo1-Slp1 interaction and inhibits APC/C activity in proliferating cells. **C**, Sgo1 bound PP2A plays a crucial role in preventing separase-mediated cleavage of Rec8, but Sgo1 does not inhibit the APC/C during meiosis.

In this study, we analyzed the ectopic expression of fission yeast Sgo1 in proliferating cells and revealed that Sgo1 holds a potential preventing cohesin cleavage during mitosis (Fig. 6B). Of note, it has been reported that budding yeast Sgo1, which when overexpressed during mitosis, prevents cohesin cleavage with cooperating with PP2A-B55 rather than PP2A-B56 (Clift et al. 2009; Yaakov et al. 2012). Because overexpression of Sgo1 in budding yeast delays securin degradation (Clift et al. 2009), so although not tested, it is possible that budding yeast Sgo1 also inhibits APC/C activity through the Sgo1 D-box. Although shugoshin has multiple faces to protect cohesin in eukaryotes, it remains an intriguing question whether the new face, which inhibits APC/C, is preserved in other organisms, including humans.

Our analysis also showed that cohesin protection during meiosis, in contrast to mitosis, is highly dependent on Sgo1-bound PP2A and not on the D-box-dependent APC/C-separase inactivation pathway (Fig. 6B). Furthermore, the D-box mutation increased Sgo1 protein abundance and protective capacity, revealing that Sgo1 protein homeostasis is dynamically regulated by the APC/C during meiosis. Although mammalian SGO2 holds the potential to inhibit separase in somatic cells (Hellmuth et al. 2020), SGO2 does not have the ability to inhibit separase in oocytes (Wetherall et al. 2025). Thus, as in fission yeast, SGO2-bound PP2A may play a major role in protecting REC8 cohesin during meiosis in higher eukaryotes. It has been shown that the reduction of REC8 cohesin and its protector SGO2 at centromeres may be a major driver of aneuploidy in aged oocytes (Chiang et al. 2010; Lister et al. 2010; Jessberger 2012; Sakakibara et al. 2015; Mihalas et al. 2023). We anticipate that research on fission yeast Sgo1 will contribute to our understanding of human SGO2 and its regulatory mechanisms, and ultimately lead to the development of methods to control aneuploidy in human oocytes.

## Materials and methods

### Schizosaccharomyces pombe strains and media

Unless otherwise stated, all media and growth conditions were as described previously (Moreno et al. 1991). *S. pombe* strains used in this study are described in Supplemental table S1. Complete medium (YE), minimal medium (SD and EMM) and sporulation media (SPA) were used. The deletion of *sgo1*^*+*^, *moa1*^*+*^, *rec12*^*+*^, *mad2*^*+*^, *par1*^*+*^, *par2*^*+*^; tagging of *sgo1*^*+*^, *par1*^*+*^, *sad1*^*+*^,*cut2*^*+*^by GFP and tagging of *sgo1*^*+*^ by mCherry were carried out according to the PCR-based gene targeting method for *S. pombe* using the *kanMX6 (kanR), hphMX6 (hygR), bsdMX6(bsdR)* and *natMX6 (natR*) genes as selection markers. The tagging of *rec8*^*+*^ by HA was performed by PCR-based genes targeting method for *S. pombe* using *ura4*^*+*^ gene as selection markers. To introduce mutations into *sgo1*^*+*^, we cloned *sgo1*^*+*^ into plasmids, and mutagenized using whole plasmid mutagenesis. Correct mutagenesis was confirmed by PCR and/or DNA sequencing. To express mCherry-Tubulin, a sequence encoding mCherry was fused to the N terminus of *atb2*^*+*^, cloned under the promoter P*adh13* (a weak version of the *adh1*^*+*^ promoter), and integrated into the locus adjacent to the *zfs1*^*+*^ gene of chromosome II (denoted the *z* locus) using the *natR* marker (Sakuno et al. 2009).

### Synchronous cultures of fission yeast meiotic cells

For meiosis microscopic observation, logarithmically growing cells were collected and resuspended in 20 mg/mL Leucine, spotted on SPA and incubated at 28°C. For chromosome segregation assay, *imr1*::GFP was observed in the *mes1*^*+*^ or *mes1-B44* mutant that arrests at prophase II (after 16 h).

### Serial dilution analyses

Thiamine was dissolved in double distilled water as a stock solution at 10 mg/ml and stored at -20°C. Sterilized EMM agar medium (115°C, 20 min) was cooled down (to about 60°C) and thiamine was added (final concentration: 5 μg/mL) to prepare the plates. For serial dilution plating assays, 5-fold dilutions of a mid-log phase culture (OD660 = 0.3∼0.6) were plated on the indicated plate and grown at 30°C; *cut1-206* cells were grown at 28°C.

### Fluorescence microscopy

All cell fluorescence microscopy was performed using a Nikon ECLIPSE Ti2-E inverted microscope with a Photometrics PRIME 95B camera. This microscope was controlled by NIS-Elements software. Nine z sections (spaced by 0.4 μm each) of the fluorescent images were converted into a single two-dimensional image by Extended Depth of Focued (EDF) module. Image J software (NIH) was used to adjust brightness and contrast.

### Time-lapse imaging

Live-cell recordings were performed at 25°C using a Nikon ECLIPSE Ti2-E inverted microscope with Nikon Intensilight C-HGFIE Precentered Fiber Illuminator, Plan-Apo 100×/1.4 oil objective. Cells were cultured in YE medium at 28°C to late log phase, spotted on SPA and incubated at 28°C for 10–11 h. Zygotes were resuspended in EMM-N were sonicated briefly and mounted on a glass-bottom dish (Bioland) coated with 500 μg/mL lectin (Sigma). Images were acquired by Z sections and create Extended Depth of Focused (EDF) document.

### Quantification of fluorescent signals

To quantify the fluorescence signal at the centromere, fluorescence images were obtained as above mentioned. Nine z sections of Sgo1-GFP signals were convert into a single two-dimensional image by Extended Depth of Focued (EDF) module. We measured the integrated intensity using a circle encompassing all dots at the centromere and subtracted the cytoplasmic signal with the same area.

### Two-hybrid assays

The yeast two-hybrid system was used according to Clontech’s instructions. We amplified the ORFs of the *slp1*^+^ and *par1*^+^ genes by PCR and cloned them into pGBK-T7, a Gal4 DNA-binding domain–based bait vector, *sgo1*^*+*^, *sgo1-N29I, sgo1-L185A, sgo1-T189A and sgo1-T189E* were cloned into pGAD-T7, a Gal4 activation domain–based prey vector. Y2HGold and Y187 strains were used for transformation of pGBK-T7 and pGAD-T7, respectively. SD-tryptophan and SD-leucine plates were used as selective medium. Positive transformants of Y2HGold and Y187 were mixed on a YPDA plate to get diploid cells and streaked on SD-Trp-Leu plate to get single colonies; then checked on nutrition-restricted plates (SD-Trp-Leu-His, SD-Trp-Leu -Ade and SD-Trp-Leu-His-Ade).

### Measurement of prometaphase duration in meiosis I

Zygotes spotted on SPA plates at 28°C for 10–11 h were used for the measurement of the prometaphase duration in meiosis I. The intervals between recordings were 1 min.

The time from Sad1-GFP separation to Cut2-GFP disappearance was measured.

### Statistical analysis

All the data replicates were applied and analyzed using GraphPad Prism version 9.5.1 (GraphPad Software). To estimate the significant differences, we used one-way ANOVA with post-hoc Tukey’s multiple comparison test.

## Acknowledgements

We thank Silke Hauf for critically reading the manuscript, Jian Chen and Jingwen Zhou for general support, and the Yeast Genetic Resource Center (YGRC) for yeast strains and plasmids. This work was supported by the National Key Research and Development Program of China (2017YFC1600403), the National Natural Science Foundation of China (Key Program, 31830068), the National Key Research and Development Program of China (2023YFF1103701) and the Entrepreneurial and innovative talent of Jiangsu (JSSCRC2021495).

## Author contributions

Ke Zhang: investigation, data curation, methodology, formal analysis. Shuchen Guo: methodology. Li Sun: supervision, methodology. Haitong Hou: validation, funding acquisition. Yoshinori Watanabe: conceptualization, data curation, formal analysis, supervision, funding acquisition, validation, investigation, methodology, and writing—original draft.

---

**Figure EV1. Related to Fig. 1.**
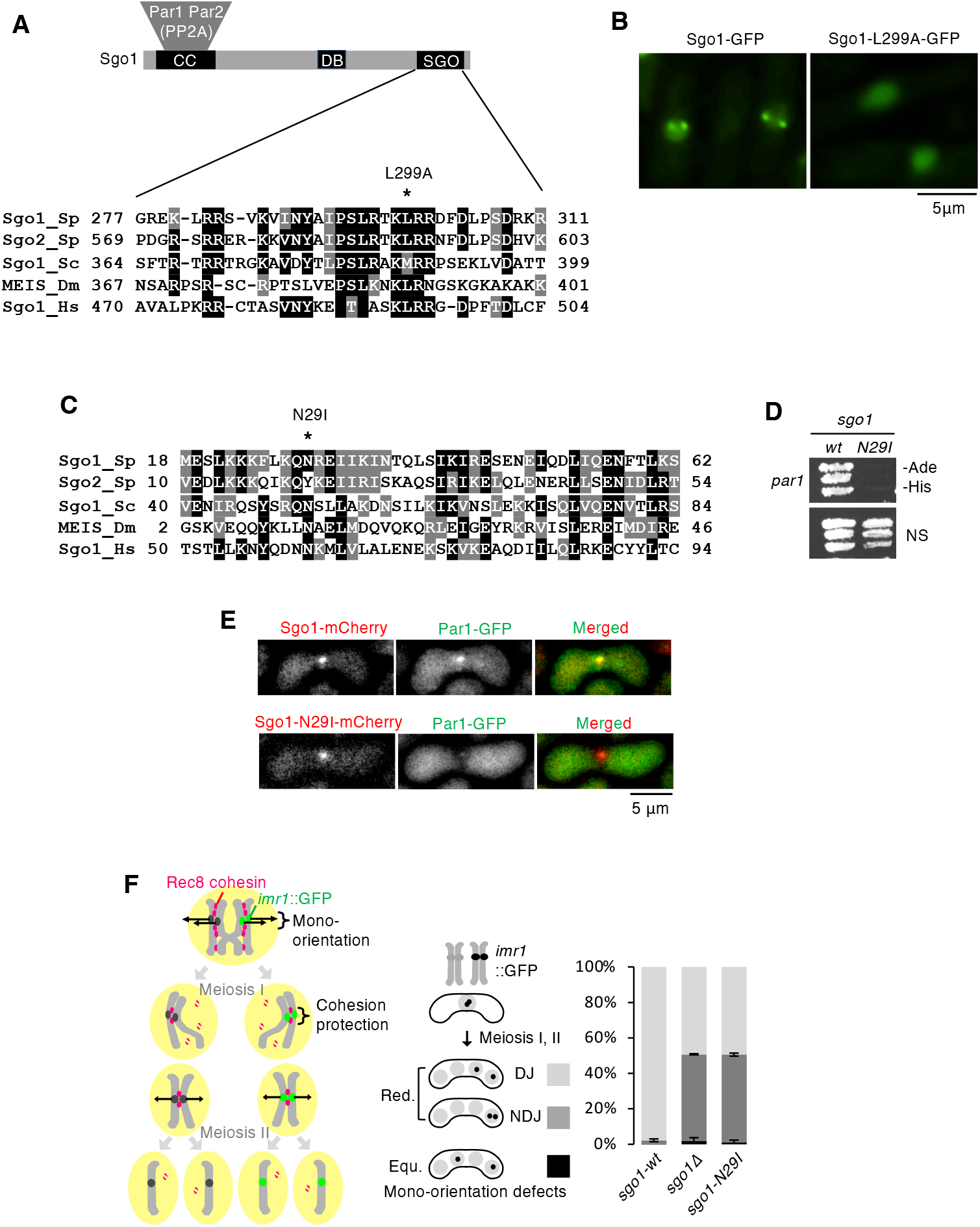
Related to Fig. 1. **A**, Alignment of the SGO motif in shugoshin family proteins of *S. pombe, S. cerevisiae, D. melanogaster* and *H. sapiens*. The mutation site L299 of *S. pombe* Sgo1 is shown. **B**, Representative fluorescent images of the indicated mitotic cells expressing Sgo1-GFP from the P*nmt1* promoter at an endogenous locus. **C**, Alignment of the coiled-coil sequence in shugoshin family proteins of *S. pombe, S. cerevisiae, D. melanogaster* and *H. sapiens*. The mutation site N29 of *S. pombe* Sgo1 is shown. **D**, Yeast two-hybrid assay testing the interaction of Sgo1 and Par1. The transformants were grown on a nonselective plate and a plate lacking adenine and histidine. **E**, Representative fluorescent images of meiotic prophase I cells expressing Par1-GFP and Sgo1-mCherryor or Sgo1-N29I-mCherry. **F**, Segregation pattern of *imr1*::GFP marked on one homolog in the indicated cells was monitored in cells after the meiosis II division. DJ: disjunction, meaning proper sister chromatid segregation in meiosis II. NDJ: nondisjunction, meaning sister chromatids move to the same pole in meiosis II, the defect originated from loss of cohesion during anaphase I. Red. (reductional): sister chromatids moved to the same pole in meiosis I. Equ. (equational): sister chromatids moved to opposite poles already in meiosis I. Error bars represent SD (three independent experiments, n > 150 cells in each experiment).

**Figure EV2. Related to Fig. 2.**
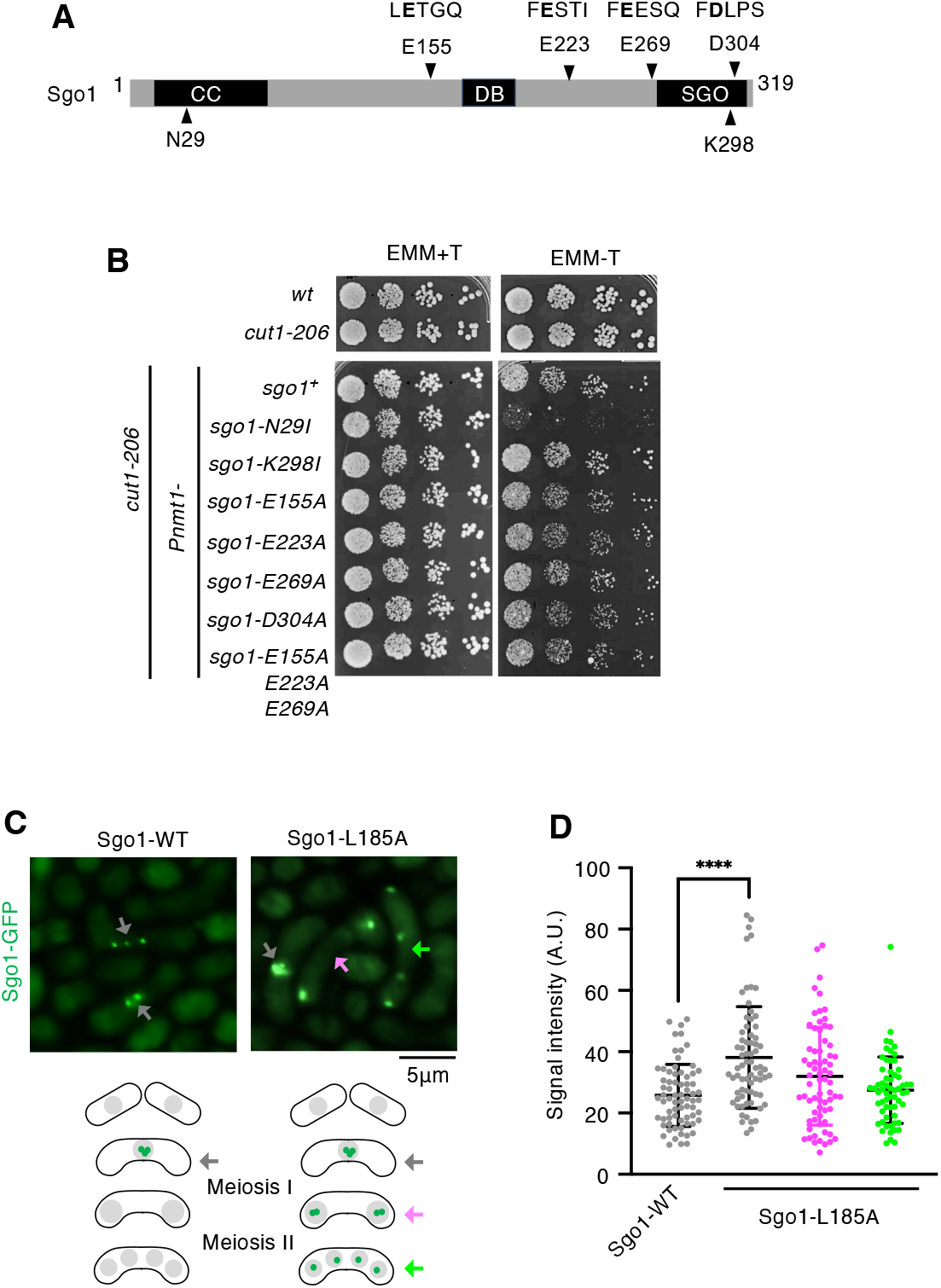
Related to Fig. 2. **A**, Schematic diagram of the Sgo1 protein showing the PP2A-interacting coiled-coil region (CC), D-box (DB), and SGO motif (SGO). Four putative pseudo-substrate sequences of Sgo1 are indicated. PP2A-binding defective mutant, N29I; centromere-localization defective mutant, K298I. **B**, Serial 5-fold dilutions of the indicated cells were spotted on EMM plates including (+T) or lacking (-T) thiamine and grown at 28C. **C**, Representative fluorescent images of the indicated meiotic cells. Schematic diagram of Sgo1-GFP localization during meiosis. GFP signals in prophase I (grey), prophase II (purple) and telophase II (green) are indicated by arrows. **D**, Quantification of Sgo1-GFP signals in prophase I (grey) in *sgo1-GFP* and *sgo1-L185A-GFP* cells as well as prophase II (purple) and telophase II (green) in *sgo1-L185A-GFP* cells. n >50 centromeres. Error bars, mean ± S.D. \*\*\*\**p* < 0.0001.

**Figure EV3. Related to Fig. 5.**
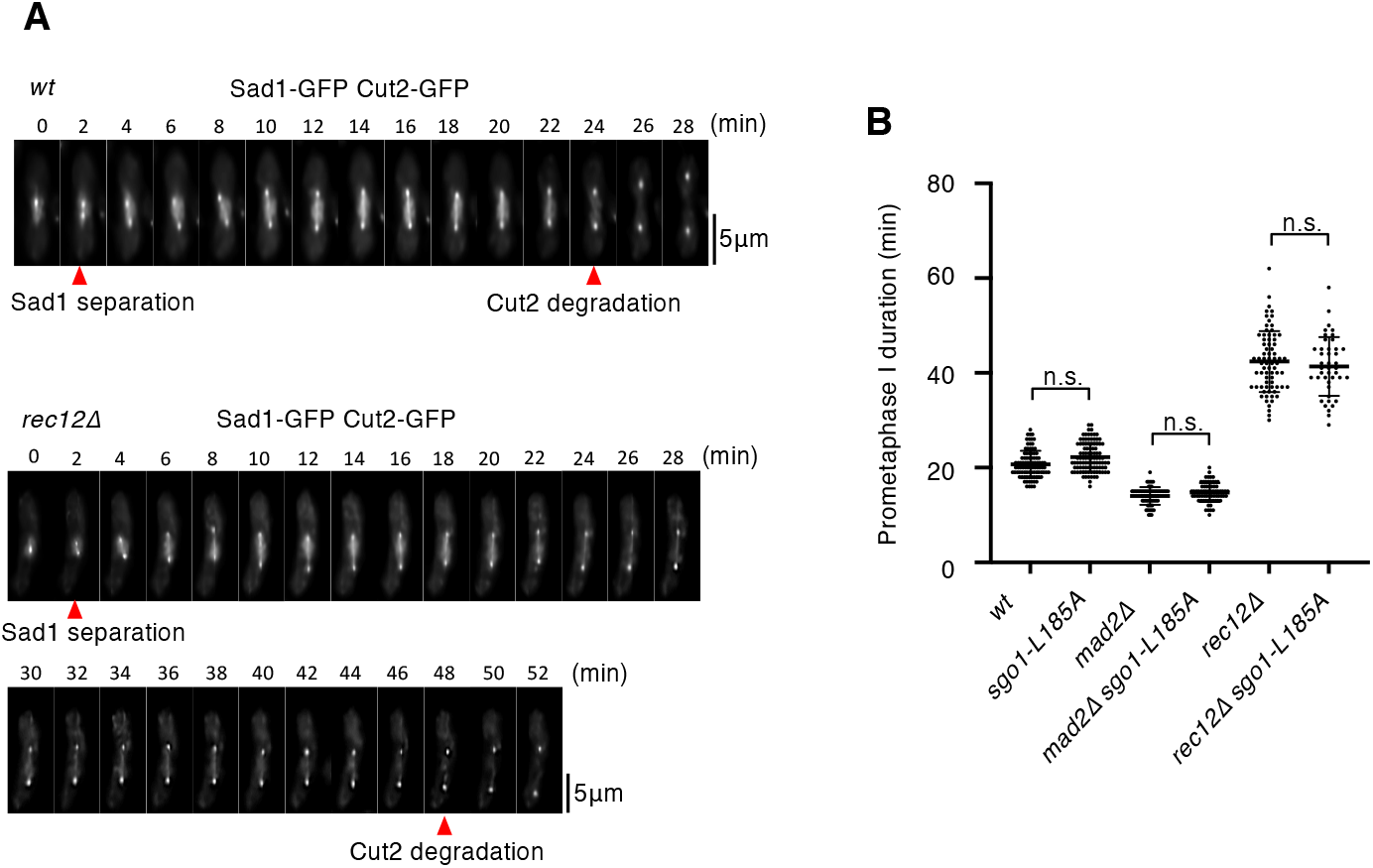
Related to Fig. 5. **A**, Cells expressing Cut2-GFP and Sad1-GFP from their endogenous loci were observed by time-lapse imaging during meiosis I at 2 min intervals. Representative timelapse images of wild-type and *rec12Δ* cells. **B**, The time from Sad1-GFP separation to Cut2-GFP disappearance of the indicated strains was measured. n > 40 cells. Each dot represents the measurement from one cell. n.s., not significant.

**Table S1.** Strain list of yeast

| Table S1 |  | Strain list of yeast |
| --- | --- | --- |
| Fig 1A |  | *h- wt*<br>*h- Pnmt41-sgo1-FLAG-GFP<<kanr*<br>*h- Pnmt1-sgo1-FLAG-GFP<<kanr*<br>*h- Padh1-rec8-3HA<<ura4+*<br>*h- Padh1-rec8-3HA<<ura4+ Pnmt41-sgo1-FLAG-GFP<<kanr*<br>*h- Padh1-rec8-3HA<<ura4+ Pnmt1-sgo1-FLAG-GFP<<kanr* |
| Fig 1B |  | *h- cut1-206*<br>*h- cut1-206 Pnmt41-sgo1-FLAG-GFP<<kanr*<br>*h- cut1-206 Pnmt1-sgo1-FLAG-GFP<<kanr*<br>*h- cut1-206*<br>*h- cut1-206 Pnmt1-sgo1-FLAG-GFP<<kanr*<br>*h- cut1-206 Pnmt1-sgo1-L299A-FLAG-GFP<<kanr* |
| Fig 1C |  | *h- wt*<br>*h- Pnmt1-sgo1-FLAG-GFP<<kanr*<br>*h- Pnmt1-sgo1-N29I-FLAG-GFP<<kanr*<br>*h- cut1-206*<br>*h- cut1-206 Pnmt1-sgo1-FLAG-GFP<<kanr*<br>*h- cut1-206 Pnmt1-sgo1-N29I-FLAG-GFP<<kanr* |
| Fig 1D |  | *h- Z::Padh13-mCherry-atb2+<<natr*<br>*h- cut1-206 Z::Padh13-mCherry-atb2+<<natr*<br>*h- cut1-206 Pnmt1-sgo1-FLAG-GFP<<kanr Z::Padh13-mCherry-atb2+<<natr*<br>*h- cut1-206 Pnmt1-sgo1-N29I-FLAG-GFP<<kanr Z::Padh13-mCherry-atb2+<<natr* |
| Fig 2B | Y2HGold | MATa, *trp1-901*, *leu2-3*, 112,*ura3-52*, *his3-200*, *gal4Δ*, *gal80Δ*,LYS2 : : GAL1<sub>UAS</sub>–<br>Gal1<sub>TATA</sub>–His3,GAL2<sub>UAS</sub>–Gal2<sub>TATA</sub>–Ade2URA3 : : MEL1<sub>UAS</sub>–Mel1<sub>TATA</sub>AUR1-C MEL1 |
|  | Y187 | MATα, *ura3-52*, *his3-200*,ade2-101, *trp1-901*, *leu2-3*, 112,*gal4Δ*, *gal80Δ*, *met–*,URA3 : :<br>GAL1<sub>UAS</sub>–Gal1<sub>TATA</sub>–LacZ,MEL1 |
| Fig 2C |  | *h- cut1-206*<br>*h- cut1-206 Pnmt1-sgo1-FLAG-GFP<<kanr*<br>*h- cut1-206 Pnmt1-sgo1-L185A-FLAG-GFP<<kanr* |
| Fig 2D |  | *h- wt*<br>*h- Pnmt1-sgo1-FLAG-GFP<<kanr*<br>*h- Pnmt1-mad2<<bsdr*<br>*h- Pnmt1-sgo1-FLAG-GFP<<kanr Pnmt1-mad2<<bsdr*<br>*h- Pnmt1-sgo1-L185A-FLAG-GFP<<kanr Pnmt1-mad2<<bsdr* |
| Fig 2E |  | *h- Pnmt1-sgo1-FLAG-GFP<<kanr Z::Padh13-mCherry-atb2+<<natr*<br>*h- Pnmt1-mad2<<bsdr Z::Padh13-mCherry-atb2+<<natr*<br>*h- Pnmt1-sgo1-FLAG-GFP<<kanr Pnmt1-mad2<<bsdr Z::Padh13-mCherry-atb2+<<natr*<br>*h- Pnmt1-sgo1-L185A-FLAG-GFP<<kanr Pnmt1-mad2<<bsdr*<br>*Z::Padh13-mCherry-atb2+<<natr* |
| Fig 3B |  | *h- cut1-206*<br>*h- cut1-206 Pnmt1-sgo1-FLAG-GFP<<kanr* |
|  |  | *h- cut1-206 Pnmt1-sgo1-T189A-FLAG-GFP<<kanr* |
|  |  | *h- cut1-206 Pnmt1-sgo1-T189E-FLAG-GFP<<kanr* |
| Fig 3C | Y2HGold | *MATa, trp1-901, leu2-3, 112,ura3-52, his3-200, gal4Δ, gal80Δ,LYS2 : : GAL1<sub>UAS</sub>Gal1<sub>TATA</sub>-His3,GAL2<sub>UAS</sub>-Gal2<sub>TATA</sub>-Ade2URA3 : : MEL1<sub>UAS</sub>-Mel1<sub>TATA</sub>AUR1-C MEL1* |
|  | Y187 | *MATα, ura3-52, his3-200,ade2-101, trp1-901, leu2-3, 112,gall4Δ, gal80Δ, met-,URA3 : : GAL1<sub>UAS</sub>-Gal1<sub>TATA</sub>-LacZ,MEL1* |
| Fig 4B |  | *h- leu1 imr1<<lacO<<ura4+ his7+<<Pdis1-lacI-GFP sgo1+<<natr* |
|  |  | *h+ sgo1+<<natr* |
|  |  | *h- leu1 imr1<<lacO<<ura4+ his7+<<Pdis1-lacI-GFP Δsgo1::natr* |
|  |  | *h+ Δsgo1::natr* |
|  |  | *h- leu1 imr1<<lacO<<ura4+ his7+<<Pdis1-lacI-GFP Δpar1::hygr* |
|  |  | *h+ Δpar1::hygr* |
|  |  | *h- leu1 imr1<<lacO<<ura4+ his7+<<Pdis1-lacI-GFP sgo1-N29I<<natr* |
|  |  | *h+ sgo1-N29I<<natr* |
|  |  | *h- leu1 imr1<<lacO<<ura4+ his7+<<Pdis1-lacI-GFP sgo1-L185A<<natr* |
|  |  | *h+ sgo1-L185A<<natr* |
|  |  | *h- leu1 imr1<<lacO<<ura4+ his7+<<Pdis1-lacI-GFP sgo1-L299A<<natr* |
|  |  | *h+ sgo1-L299A<<natr* |
| Fig 4C |  | *h- Pnmt1-sgo1-FLAG-GFP<<kanr* |
|  |  | *h- Pnmt1-sgo1-FLAG-GFP<<kanr Δpar1::hygr Δpar2::natr* |
|  |  | *h- Pnmt1-sgo1-N29I-FLAG-GFP<<kanr* |
|  |  | *h- Pnmt1-sgo1-L185A-FLAG-GFP<<kanr* |
|  |  | *h- Pnmt1-sgo1-L299A-FLAG-GFP<<kanr* |
|  |  | *h- Padh1-rec8-3HA<<ura4+ Pnmt1-sgo1-FLAG-GFP<<kanr* |
|  |  | *h- Padh1-rec8-3HA<<ura4+ Pnmt1-sgo1-FLAG-GFP<<kanr Δpar1::hygr Δpar2::natr* |
|  |  | *h- Padh1-rec8-3HA<<ura4+ Pnmt1-sgo1-N29I-FLAG-GFP<<kanr* |
|  |  | *h- Padh1-rec8-3HA<<ura4+ Pnmt1-sgo1-L185A-FLAG-GFP<<kanr* |
|  |  | *h- Padh1-rec8-3HA<<ura4+ Pnmt1-sgo1-L299A-FLAG-GFP<<kanr* |
| Fig5A |  | *h90 imr1<<lacO<<ura4+ his7+<<Pdis1-lacI-GFP mes1-B44 cut1-206 sgo1+<<natr* |
| Fig5B |  | *h90 imr1<<lacO<<ura4+ his7+<<Pdis1-lacI-GFP mes1-B44 cut1-206 sgo1+<<natr* |
|  |  | *h90 imr1<<lacO<<ura4+ his7+<<Pdis1-lacI-GFP mes1-B44 cut1-206 sgo1-L185A<<natr* |
|  |  | *h90 imr1<<lacO<<ura4+ his7+<<Pdis1-lacI-GFP mes1-B44 cut1-206 sgo1-N29I L185A<<natr* |
|  |  | *h90 imr1<<lacO<<ura4+ his7+<<Pdis1-lacI-GFP mes1-B44 cut1-206 Δsgo1::natr* |
| Fig5C,D |  | *h- leu1 imr1<<lacO<<ura4+ his7+<<Pdis1-lacI-GFP* |
|  |  | *h+ wt* |
|  |  | *h- leu1 imr1<<lacO<<ura4+ his7+<<Pdis1-lacI-GFP Δmoa1::natr* |
|  |  | *h+ Δmoa1::natr* |
|  |  | *h- leu1 imr1<<lacO<<ura4+ his7+<<Pdis1-lacI-GFP sgo1-L185A<<hygr* |
|  |  | *h+ sgo1-L185A<<hygr* |
|  |  | *h- leu1 imr1<<lacO<<ura4+ his7+<<Pdis1-lacI-GFP Δmoa1::kanr sgo1-L185A<<hygr* |
|  |  | *h+ Δmoa1::kanr sgo1-L185A<<hygr* |
| FigEV1 B |  | *h- Pnmt1-sgo1-FLAG-GFP<<kanr* |
|  |  | *h- Pnmt1-sgo1-L299A-FLAG-GFP<<kanr* |
| FigEV1 D | Y2HGold | *MATa, trp1-901, leu2-3, 112,ura3-52, his3-200, gal4Δ, gal80Δ,LYS2 : : GAL1<sub>UAS</sub>–Gal1<sub>TATA</sub>–His3,GAL2<sub>UAS</sub>–Gal2<sub>TATA</sub>–Ade2URA3 : : MEL1<sub>UAS</sub>–Mel1<sub>TATA</sub>AUR1-C MEL1* |
|  | Y187 | *MATα, ura3-52, his3-200,ade2-101, trp1-901, leu2-3, 112,gall4Δ, gal80Δ, met–,URA3 : : GAL1<sub>UAS</sub>–Gal1<sub>TATA</sub>–LacZ,MEL1* |
| FigEV1 E |  | *h90 sgo1-mCherry<<natr par1-GFP<<kanr* |
|  |  | *h90 sgo1-N29I-mCherry<<natr par1-GFP<<kanr* |
| FigEV1 F |  | *h- leu1 imr1<<lacO<<ura4+ his7+<<Pdis1-lacI-GFP sgo1-GFP<<natr* |
|  |  | *h+ sgo1-GFP<<natr* |
|  |  | *h- leu1 imr1<<lacO<<ura4+ his7+<<Pdis1-lacI-GFP Δsgo1::natr* |
|  |  | *h+ Δsgo1::natr* |
|  |  | *h- leu1 imr1<<lacO<<ura4+ his7+<<Pdis1-lacI-GFP sgo1-N29I-GFP<<natr* |
|  |  | *h+ sgo1-N29I-GFP<<natr* |
| FigEV2 B |  | *h- wt* |
|  |  | *h- cut1-206* |
|  |  | *h- cut1-206 Pnmt1-sgo1-FLAG-GFP<<kanr* |
|  |  | *h- cut1-206 Pnmt1-sgo1-N29I-FLAG-GFP<<kanr* |
|  |  | *h- cut1-206 Pnmt1-sgo1-K298I-FLAG-GFP<<kanr* |
|  |  | *h- cut1-206 Pnmt1-sgo1-E155A-FLAG-GFP<<kanr* |
|  |  | *h- cut1-206 Pnmt1-sgo1-E223A-FLAG-GFP<<kanr* |
|  |  | *h- cut1-206 Pnmt1-sgo1-E269A-FLAG-GFP<<kanr* |
|  |  | *h- cut1-206 Pnmt1-sgo1-D304A-FLAG-GFP<<kanr* |
|  |  | *h- cut1-206 Pnmt1-sgo1-E155AE223AE269A-FLAG-GFP<<kanr* |
| FigEV2 C,D |  | *h90 sgo1-GFP<<natr* |
|  |  | *h90 sgo1-L185A-GFP<<natr* |
| FigEV3 A |  | *h90 sad1-GFP<<hygr cut2-GFP<<kanr sgo1+<<natr* |
|  |  | *h90 sad1-GFP<<hygr cut2-GFP<<kanr Δrec12::bsdr* |
| FigEV3 B |  | *h90 sad1-GFP<<hygr cut2-GFP<<kanr sgo1+<<natr* |
|  |  | *h90 sad1-GFP<<hygr cut2-GFP<<kanr sgo1-L185A<<natr* |
|  |  | *h90 sad1-GFP<<hygr cut2-GFP<<kanr Δmad2::natr* |
|  |  | *h90 sad1-GFP<<hygr cut2-GFP<<kanr Δmad2::bsdr sgo1-L185A<<natr* |
|  |  | *h90 sad1-GFP<<hygr cut2-GFP<<kanr Δrec12::bsdr* |
|  |  | *h90 sad1-GFP<<hygr cut2-GFP<<kanr Δrec12::bsdr sgo1-L185A<<natr* |

